# Loss of white matter tracts and persistent microglial activation in the chronic phase of ischemic stroke in female rats and the effect of miR-20a-3p treatment

**DOI:** 10.1101/2025.02.01.636074

**Authors:** Dayalan Sampath, Macy E. Zardeneta, Zara Akbari, Jacob Singer, Balaji Gopalakrishnan, David Austin Hurst, Monserrat Villarreal, Erin A. McDaniel, Brenda Patricia Noarbe, Andre Obenaus, Farida Sohrabji

## Abstract

Our previous studies showed that intravenous injections of the small non-coding RNA mir-20a-3p is neuroprotective for stroke in the acute phase and attenuates long-term cognitive impairment in middle-aged female rats. In this study, we evaluated postmortem brain pathology at 100+d after stroke in a set of behaviorally characterized animals. This included Sham (no stroke) controls or stroke animals that received either mir20a-3p at 4h, 24h and 70d iv post stroke (MCAo+mir20a-3p) or a scrambled oligo (MCAo+Scr). Brain volumetric features were analyzed with T2 weighted and Diffusion Tensor magnetic resonance imaging (MRI) followed by histological analysis. Principal component analysis of Fractional Anisotropy (FA)-diffusion tensor MRI measures showed that MCAo+Scr and MCAo+mir20a-3p groups differed significantly in the volume of white matter but not gray matter. Weil myelin-stained sections confirmed decreased volume of the corpus callosum, internal capsule and the anterior commissure in the ischemic hemisphere of MCAo+Scr animals compared to the non-ischemic hemisphere, while sham and MCAo+Mir-20a-3p showed no hemispheric asymmetries. The MCAo+Scr group also exhibited asymmetry in hemisphere and lateral ventricle volumes, with ventricular enlargement in the ischemic hemisphere as compared to the non-ischemic hemisphere. The numbers of microglia were significantly elevated in white matter tracts in the MCAo+Scr group, with a trend towards increased myelin phagocytic microglia in these tracts. Regression analysis indicated that performance on an episodic memory test (novel object recognition test; NORT) was associated with decreased white matter volume and increased microglial numbers. These data support the hypothesis that stroke-induced cognitive impairment is accompanied by white matter attrition and persistent microglial activation and is consistent with reports that cognitive deterioration resulting from vascular diseases, such as stroke, is associated with secondary neurodegeneration in regions distal from the initial infarction.

## INTRODUCTION

Stroke is the fifth leading cause of death in the United States, with a higher incidence of severity among middle-aged women (55-64 years) than age-matched men ^1,2, 3,4^. Stroke patients have a 9-fold increased risk of developing mild cognitive impairment ^5^ and 4-fold increased risk of dementia compared to non-stroke subjects ^6^. Recent studies have reported that women have worse cognitive outcomes in the early post stroke period (90 days after stroke) compared to men, which was attributable to sociodemographic and pre-stroke characteristics, especially widowhood status ^7, 8^. Interestingly, a large multicenter study of stroke patients reported that while post-stroke cognitive impairment was similar in males and females (51%), sex differences were noted in the domains of cognitive impairment, with women exhibiting great impairments in attention, executive functioning and language, while men had higher risk of verbal memory impairment ^9^.

In the acute phase of stroke, neuronal death is rapid, primarily caused by excitotoxicity, oxidative stress, and neuroinflammation^10^. Subsequently, in the chronic phase (weeks to months), structural and functional alterations continue to occur in regions distal to the ischemic zone, a phenomenon termed connectional diaschisis^11^, and includes inadequate remyelination^12^ and extensive functional reorganization^13^. Loss of white matter is a known long-term effect of ischemic stroke, resulting in inefficient information transfer throughout the brain^14,15^.

Microglia play an important role during both acute and chronic phases after ischemia. In response to injury signals, resting microglia become activated and motile, migrating to the ischemic zone to phagocytize cellular debris. In the chronic phase, microglia can remain activated and this persistent activation is seen in regions distal from the ischemic zone ^16^. In an embolic stroke model, neuroinflammation was virtually undetectable near the infarction at 7 months post-stroke while histological and T2 MRI analysis showed microglial activation in the thalamus, a region remote from the ischemic location^17^. In an animal model of diabetes, modulation of microglial polarization prevented the development of post-stroke cognitive impairment ^18^ while knockdown of microglia, by silencing colony stimulating factor 1 receptor (CSF1R), attenuated long-term inflammation, increased white matter myelination after stroke and prevented cognitive decline^19^. Collectively these studies indicate that chronic microglial activation may be associated with delayed neurodegeneration and post-stroke cognitive impairment.

Our previous studies have shown that in middle-aged acyclic female rats miR-20a-3p, a small non-coding RNA, reduces infarct volume and sensorimotor deficits acutely ^20^ and attenuates cognitive decline in the chronic phase (1-3 months)^21^. In the present study, we evaluated long term changes in histomorphometry due to stroke and mir-20a-3p treatment. A combination of diffusion tensor magnetic resonance imaging (DT-MRI), histology and immunohistochemistry indicate that ischemic stroke resulted in damage to white matter tracts, with evidence of phagocytic microglia when measured 3+ months after stroke. Moreover, these measures were attenuated in animals treated with mir-20a-3p. Regression analysis indicated that decreased volume of forebrain white matter tracts and increased proportion of microglia were predictive of episodic memory impairment due to stroke.

## METHODS

This study used brain tissue from a behaviorally characterized cohort of middle-aged, female Sprague Dawley rats. Detailed methodology of this cohort is available in Sampath et al ^21^ and the experimental plan is shown in Supplementary Fig S1.

### Animals

Eleven-month-old female Sprague Dawley rats were purchased from Envigo (IN). Rats were subject to daily vaginal smears for 14 days to determine estrus cyclicity and only those rats that were in constant diestrus (a persistent low estrogen state) were included in the study.

### Middle cerebral artery occlusion

Animals were anesthetized using 87 mg/kg of body weight (BWT) ketamine and 10 mg/ kg of BWT xylazine. All rats were subjected to left-sided MCAo^22,23^ induced by stereotaxic injection of endothelin-1 (3 μl of 0.5 mg/ml ET-1 stock, American Peptide Company) adjacent to the middle cerebral artery (0.9 mm anterior to Bregma, 3.4 mm lateral to bregma on the left side and −8.5 mm ventral relative to the dura). Sham animals (no stroke controls) were anesthetized and placed in the stereotaxic frame, and a burr hole was drilled at the same coordinates but not injected with ET-1.

### Drug treatment

MicroRNA (miR-20a-3p; ACU GCA UUA CGA GCA CUU ACA) oligonucleotide sequence/scrambled negative control (Thermofisher Scientific, Grand Island, NY) was mixed with in vivo RNA-LANCEr II (Bioo Scientific, Austin, TX), using our previous protocols^24, 25,20,26^, and injected intravenously (tail vein) (7 μg/kg) at 4 hours, 24 hours, and 70 days after stroke.

### Acute tests of sensorimotor integrity and chronic tests of cognition

Animals were subjected to tests of sensorimotor integrity pre- and post-stroke. This included the vibrissae-evoked forelimb placement test and the adhesive removal test. Animals were tested for long-term cognitive function at 30, 60 and 100d after stroke, namely remote fear memory retrieval, and the novel object recognition test ^21^.

### Magnetic resonance imaging

#### MRI acquisition

Brain imaging was performed on postmortem tissue. At 100+d after stroke, rats were deeply anesthetized and perfused transcardially with saline, followed by 4% paraformaldehyde, and stored in 1X PBS+ 0.01% sodium azide. Post-mortem brain tissue remained inside the skull and was imaged on a high-resolution MRI scanner at the Preclinical and Translational Imaging Center at University of California Irvine (9.4 T; Bruker Bio Spin, Paravision 5.1). Diffusion tensor imaging (DTI) and T2-weighted imaging (T2WI) scans were acquired for 1 hour 52 minutes and 38 minutes, respectively. Both scans were acquired with the same spatial resolution: 1.5cm field of view, 0.5mm slice thickness, and a 128x128 acquisition matrix. DTI images were collected with repetition time/echo time (TR/TE) = 8000 msec/35.7 msec, and 30 directions (5 b=0mT/m, b=3000mT/m). T2WI scans were acquired with 10 echoes and TR/TE = 4000 msec/10 msec.

#### MRI Analysis

Experimenters were blinded to animal group assignment during MRI analysis. The skull was stripped from the MRI scan using ITK-SNAP (version 3.6.0)^27^. DTI images then underwent correction for bias field inhomogeneities ^28^ and eddy current distortions. Using FMRIB’s Diffusion Toolbox from FMRIB’s Software Library (FSL), diffusion tensor models were assigned to each voxel to generate DTI parametric maps: fractional anisotropy (FA), axial (AxD), mean (MD), radial (RD) diffusivity. T2WI bias field inhomogeneities were corrected and quantitative T2 relaxation maps were generated with JIM software (Xinapse Systems Ltd; West Bergholt, Essex; United Kingdom). One hundred and twenty four regional labels (64 bilateral regions) from the Waxholm Rat Brain Atlas ^29^ were applied to each animal using the Advanced Normalization Tools (ANTS) and regional DTI metrics were extracted. Each atlas overlay was manually inspected, and misalignments were corrected with ITK-SNAP. Regional metric outliers within each group were excluded with 1.5xIQR above the first quartile or below the third quartile including all regions containing less than 20 voxels, yielding a total of 78 bilateral regions. The list was further curated for regions associated with forebrain cognitive circuits resulting in 29 regions for further analysis.

Tractography was performed in DSI Studio software (Chen-Jan 25, 2022) and whole brain reconstruction was used. Regions of interest (ROI) for white matter tracts passing through the bilateral anterior commissure anterior (ACA), and the basolateral nucleus of amygdala (BLA) were reconstructed using a diffusion sampling length ratio of 0.75. The location of ACA was confirmed by Paxinos and Watson atlas coordinates (2.20mm anterior to bregma) as circular hyperintense regions on FA parametric maps. Deterministic tractography was performed using an angular threshold of 70, a step size of 0.06, 0.80 smoothing and 1 million seed regions were assigned to reconstruct tracts from the ACA and passing through the BLA region. The differences in tract metrics and their characteristics between the ipsilateral and contralateral hemispheres were obtained, including number of tracts, elongation, volume and total surface area; where volume = number of tracts X voxel size, elongation =length of the tract/diameter of the tract (elongation=length/diameter), 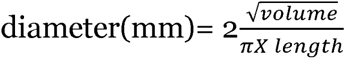, surface area= number of surface voxels X voxel spacing^30^.

#### Principal component analysis

To evaluate brain features associated with stroke and microRNA treatment, principal component analysis of DT-MRI images was performed using Metaboanalyst (www.metaboanalyst.ca). In each case, the average of the left (ischemic) and right (non-ischemic hemisphere) FA measures were used for PCA analysis. PCA analysis was first performed for all regions followed by separate analysis for gray matter and white matter. In addition, gray matter and white matter regions associated with the hippocampus were analyzed separately. Group differences in the centroid and dispersion of the PCA plots were determined by PERMANOVA and dispersion was considered significant at p≤0.05.

### Histological analysis

After MR imaging, the brains were removed from the skull and stored in 4%PFA overnight, and then transferred to a sucrose (10%) and sodium azide (0.02%) solution. Brains were shipped to Neuroscience Associates Inc (TN, USA) for block embedding. Each block contained 16 brains with brains from each group spread across each block in a randomized manner. Blocks were sectioned at 40 microns and one set of sections was stained for Weil’s myelin stain using NSA proprietary protocols.

Weil Myelin-stained sections were imaged at 20X magnification using an Olympus VS120 Slide Scanner. In addition to myelinated pathways, hemisphere and lateral ventricles were clearly delineated in Weil myelin-stained sections. For the quantification of hemispheric volumes, each hemisphere of brain sections between bregma +4.20 mm- (-) 4.52mm (23 sections) were manually outlined and measured with an open-source software QuPAth (0.4.3) to obtain the area measurement (μm^2)^. The sum of the cross-sectional areas was then multiplied by the section thickness and the number of sections measured to obtain the total volume (μm^3^). Lateral ventricle volume on each hemisphere was measured between bregma +1.60 mm- -1.80 mm (7 sections) for each brain. Corpus callosum volume was evaluated between bregma +2.70 mm- (-) 7.30 mm (25-26 sections for each brain), the internal capsule volume was quantified from bregma -0.04 mm- (-)3.6 mm (6 sections), and the anterior commissure from bregma +1.70 mm- (+) 0.20 mm Anterior – Posterior (3 sections).

### Microglia Analysis

Block-embedded brain sections through the corpus callosum and internal capsule were probed with antibodies for Ionized calcium binding adapter molecule 1 (Iba1) and degraded myelin basic protein (dMBP). Floating sections were washed three times (5 min) in TBS plus Tween and incubated in 1X antigen retrieval solution (Abcam Cat. #93678) at 80°C for 20 minutes. Sections were incubated in blocking solution (10% Normal Donkey Serum, 0.1% Triton-X100 in TBST) for 1 hr at RT, followed by incubation in 1% NDS and 0.01% Triton-X100 in TBST and primary antibodies goat anti-IBA1 (1:500; Novus Biologicals, NB100-1028); rabbit anti-dMBP (1:1000; Millipore Sigma, AB5864), for 24hr at 4°C. After washes, sections were incubated with secondary antibodies (1:1000) donkey anti-Rabbit Alexa Fluor 555 (Thermo Scientific #A-31572), and donkey anti-Goat Alexa Fluor 488 (Thermo Scientific #A-11055) for 1 hr at RT. Sections were mounted on glass slides and coverslipped with ProLong Gold Antifade mounting medium (Thermo Fisher, P36934).

#### Quantitative analysis

For each selected tract, coronal brain sections were imaged on a confocal scanning laser microscope (FV3000, Olympus) with Fluoview software. Image capture and analysis was performed by separate experimenter’s blinded to treatment conditions. For each image, all single-labeled (Iba1) and double-labeled (Iba1+dMBP) cells were counted in the left and right hemisphere. For the corpus callosum, microglia were analyzed in the genu and splenium and summed together. In both tracts, the number of Iba1+ microglia and the number of Iba1+dMBP double-labeled microglia in the ischemic hemisphere were normalized to the non-ischemic hemisphere.

### Tomato Lectin histochemistry

Block embedded brain sections were washed in PBS and blocked with 5% BSA for one hour. Free floating sections were incubated with tomato lectin (FL-1171; Vector Laboratories CA; 1:500 concentration in PBS) over night and washed 3x in PBS before mounting on glass slides. Slides were cover slipped with a glycerol-based mounting media (DABCO^TM^) Electron Microscopy Sciences Inc). Sections were then imaged in a slide scanner (Olympus VS_120_) at 20X magnification. Images were visualized in QuPath (0.4.3) software and 3 separate grids were demarcated in the cortex and striatum of each image. Within each grid, lectin-stained vessels were quantified for their density and length with Fiji (Image J2). Micro vessels were segmented from the image background with Weka Segmentation plugin for ImageJ. The segmented vessels were quantified for vessel morphometry (micro vessel density and vessel length density) with vascular plugin. Cortical and striatal vessel density was analyzed separately for each hemisphere.

### Multiple regression analysis

Multiple regression was performed using the following histological measures: normalized volume of the anterior commissure, corpus callosum, internal capsule, the number of microglia and myelin phagocytic microglia in the corpus callosum and internal capsule. The dependent variable was either performance on the NORT test or Remote Fear memory retrieval. Regression analysis using the Least Squares method was performed on GraphPad Prism software (Version 10.3.1). Confidence intervals were considered significant at p<0.05.

## RESULTS

Histomorphometry was performed on a set of behaviorally characterized female Sprague Dawley rats after stroke as reported in our recent publication^21^. Animals were tested periodically 30-100 days after stroke. Vehicle (scramble) treated MCAo animals showed persistent impairment in episodic memory (Novel Object Recognition Test) (Supplementary Fig S2A) and early loss of associative memory recall (remote fear memory retrieval) (Supplementary Fig S2B). Mir-20a-3p treatment mitigated performance in both these tests.

### Atlas mapping to MR images and principal component analysis

ROI were obtained from Fractional anisotropy (FA) of DT-MRI encompassing gray and white matter regions. Principal component analyses (PCA) was performed to detect whether these features (brain regions defined through atlas-mapping) differed among the groups due to stroke and microRNA treatment. PCA comparisons of 29 brain regions showed that centroid and dispersion was significantly different among Sham, MCAo+Scr, MCAo+Mir-20a-3p groups (F: 3.6585; R-squared: 0.19608; p: 0.01; Fig 1A). PCA comparisons of the stroke groups (MCAo+Scr, MCAo+Mir20a-3p) also showed significant dispersion (F: 5.8709; R-squared: 0.20335; p: 0.001; Fig 1B). Scree plot analysis (Fig 1C) showed that 71.4% of the variance between groups was explained by principal components 1 through 4. Features expressed in at least 2 of the components and contributing at least 15% to each of these components are shown in Table 1A. Of the 9 features thus curated, 5 were white matter tracts, 3 gray matter regions, and the ventricular system.

**Figure 1.**
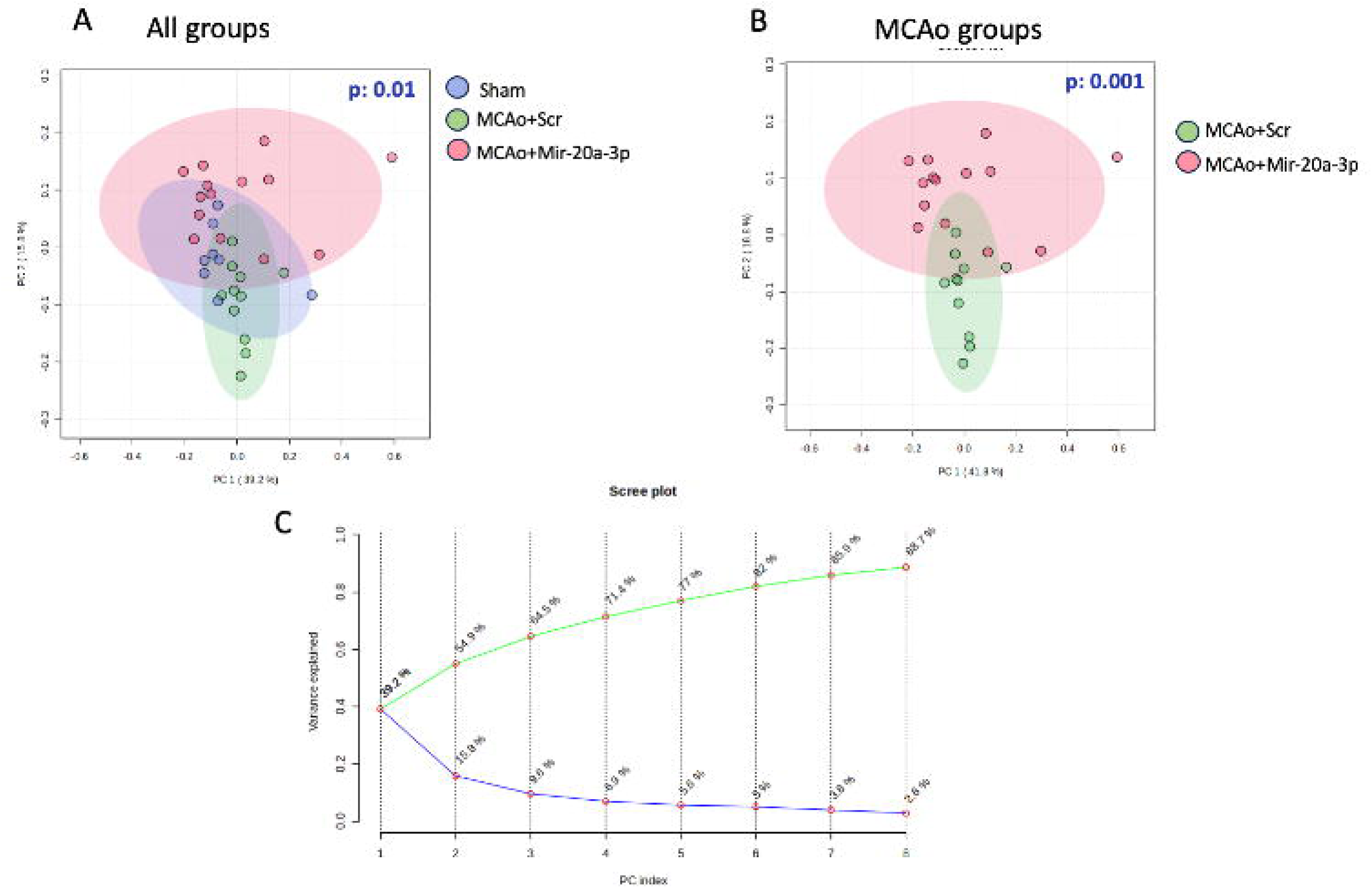
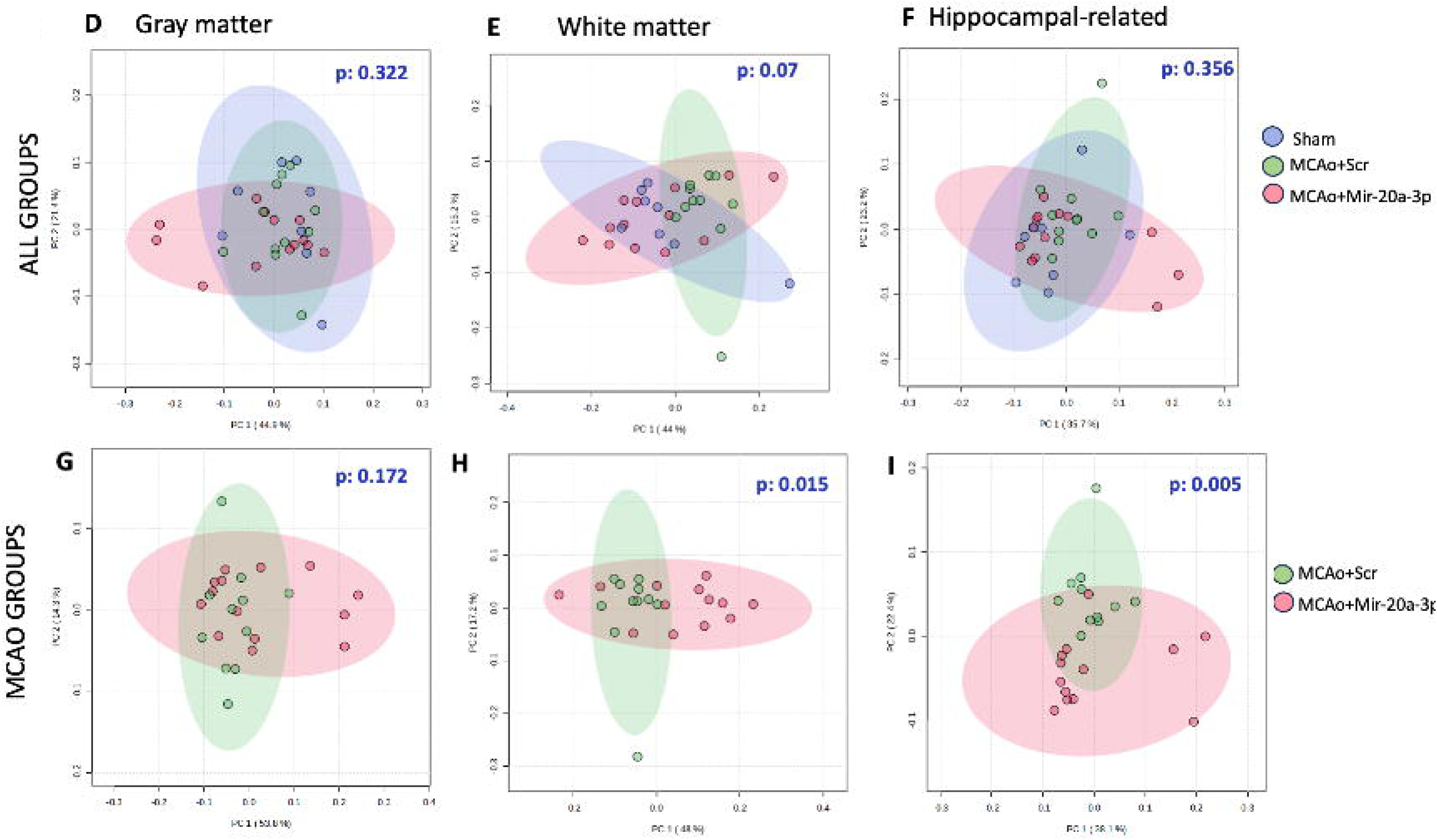
Principal Component Analysis of Fractional anisotropy-DTI measures: ROI were obtained from Fractional anisotropy (FA) of DTI MRI plots for 29 regions. For each region, ROI are averages of left and right sides. A. PCA plot comparing all groups: Sham, MCAo+Scr, and MCAo-mir20a-3p B. PCA plots comparing stroke groups only: MCAo+Scr, and MCAo-mir20a-3p. C. Scree plot loading indicated that 71.4 % of the variance between groups was attributed to regions in the first 4 principal components (detailed in Table 1A). PCA plots comparing Sham, MCAo+Scr, and MCAo-mir20a-3p groups for gray matter (D), white matter (E) and hippocampal-related features (F). PCA plots comparing MCAo+Scr, and MCAo-mir20a-3p groups for gray matter (G), white matter (H) and hippocampal-related features (I). Centroid and dispersion measures were evaluated by Permanova, exact p values are shown on the plots. N= 8 in sham, 11 in MCAo+Scrambled (MCAo+Scr), and 12 in MCAo+ miR-20a-3p groups.

**TABLE 1A.** Features from PC1, PC2, PC3, PC4 that contribute >15% variance.

| Region | Component | Classification |
| --- | --- | --- |
| Anterior_Commissure | PC1, 3 | White matter |
| Anterior_Commissure_Anterior_Part | PC1, 2, 3 | White matter |
| Cingulate_Cortex_Area_2 | PC1,2,3,4 | Gray matter |
| Corpus_Callosum_And_Associated_Subcortical_White_Matter | PC1,2,3,4 | White matter |
| Descending_Corticofugal_Pathways | PC1, 2,3 | White matter |
| Glomerular_Layer_Of_The_Olfactory_Bulb | PC1, 3 | Gray matter |
| Perirhinal_Area_35 | PC1,3 | Gray matter |
| Ventral_Hippocampal_Commissure | PC3,4 | White matter |
| Ventricular_System | PC2,4 | - |

Next, PCA comparisons were performed separately for gray matter, white matter and regions (gray and white) associated with the hippocampus (shown in Table 1B).

**TABLE 1B.** DT-MRI features included in PCA analysis of Gray matter, White matter and Hippocampal related.

| **Gray Matter** | **White Matter** | **Hippocampal related** |
| --- | --- | --- |
| Basal_Forebrain_Region | Alveus_Of_The_Hippocampus | Alveus_Of_The_Hippocampus |
| Cingulate_Cortex_Area_2 | Anterior_Commissure | Cornu_Ammonis_1 |
| Dentate_Gyrus | Anterior_Commissure_Anterior_Part | Cornu_Ammonis_2 |
| Entorhinal_Cortex | Anterior_Commissure_Posterior_Part | Cornu_Ammonis_3 |
| Frontal_Association_Cortex | Corpus_Callosum_&_Associated<br>Subcortical_White_Matter | Dentate_Gyrus |
| Neocortex | Descending_Corticofugal_Pathways | Entorhinal_Cortex |
| Perirhinal_Area_35 | Fimbria_Of_The_Hippocampus | Fimbria_Of_The_Hippocampus |
| Perirhinal_Area_36 | Fornix | Fornix |
| Striatum | Ventral_Hippocampal_Commissure | Perirhinal_Area_35 |
| Subiculum |  | Perirhinal_Area_36 |
| Substantia_Nigra |  | Subiculum |
|  |  | Ventral_Hippocampal_Commissure |

PCA comparisons for gray matter regions of all groups (Sham, MCAo+Scr, MCAo+Mir-20a-3p) showed considerable overlap in the centroid and dispersion, indicating that the groups did not differ (Fig 1D; F: 0.94087; R-squared: 0.060934; p-value: 0.475). However, there was a statistical trend toward group dispersion in white matter structures (Fig 1E; F: 2.4549; R-squared: 0.14479; p: 0.07). A combination of gray and white matter regions related to hippocampal structures were also assessed, due to the role of the hippocampus in fear retrieval memory and episodic memory tested in this cohort, but did not show significant dispersion (Fig 1F, F: 1.0904; R-squared: 0.069939; p: 0.356).

Next, shams were removed and PCA analyses was performed for the two stroke groups (MCAo+Scr, MCAo+Mir-20a-3p) to assess the impact of miR-20a-3p treatment on FA measurements after stroke. No significant dispersion was observed between the MCAo groups for features related to gray matter (Fig 1G; F: 1.1388; R-squared: 0.049216; p: 0.322). However, scrambled and mir-20a-3p treated MCAo groups were significantly different in features related to white matter (Fig 1H; F: 5.1003; R-squared: 0.1882; p: 0.015) as well as hippocampal related structures (Fig 1I; F: 5.4922; R-squared: 0.19977; p: 0.005), indicating an effect of miRNA treatment.

### Deterministic tractography: miR-20a-3p mitigates post-stroke, left-right asymmetries in white matter tracts

Whole brain tractography was used to visualize the tract architecture of major white matter tracts and their connections with neighboring gray matter areas (Fig 2i). Rostrocaudal array of coronal images at the level of the (a) neocortex (Bregma +2.70), (b) striatum (Bregma +1.00), (c) dorsal hippocampus (Bregma -3.30), (d) dorsoventral hippocampus (Bregma -4.30) and (e) cerebellum (Bregma -11.80) revealed qualitative differences in white matter density, especially in the MCAo+Scr group. For example, the cingulum bundle (white arrow; Fig 2ib), the corpus callosum (white arrow; Fig 2ic) as well as cerebellar white matter (white arrow; Fig 2ie) were less dense in the MCAo+scr group (middle panel), compared to Sham (upper panel) and MCAo+Mir-20a-3p (bottom panel).

**Fig 2:**
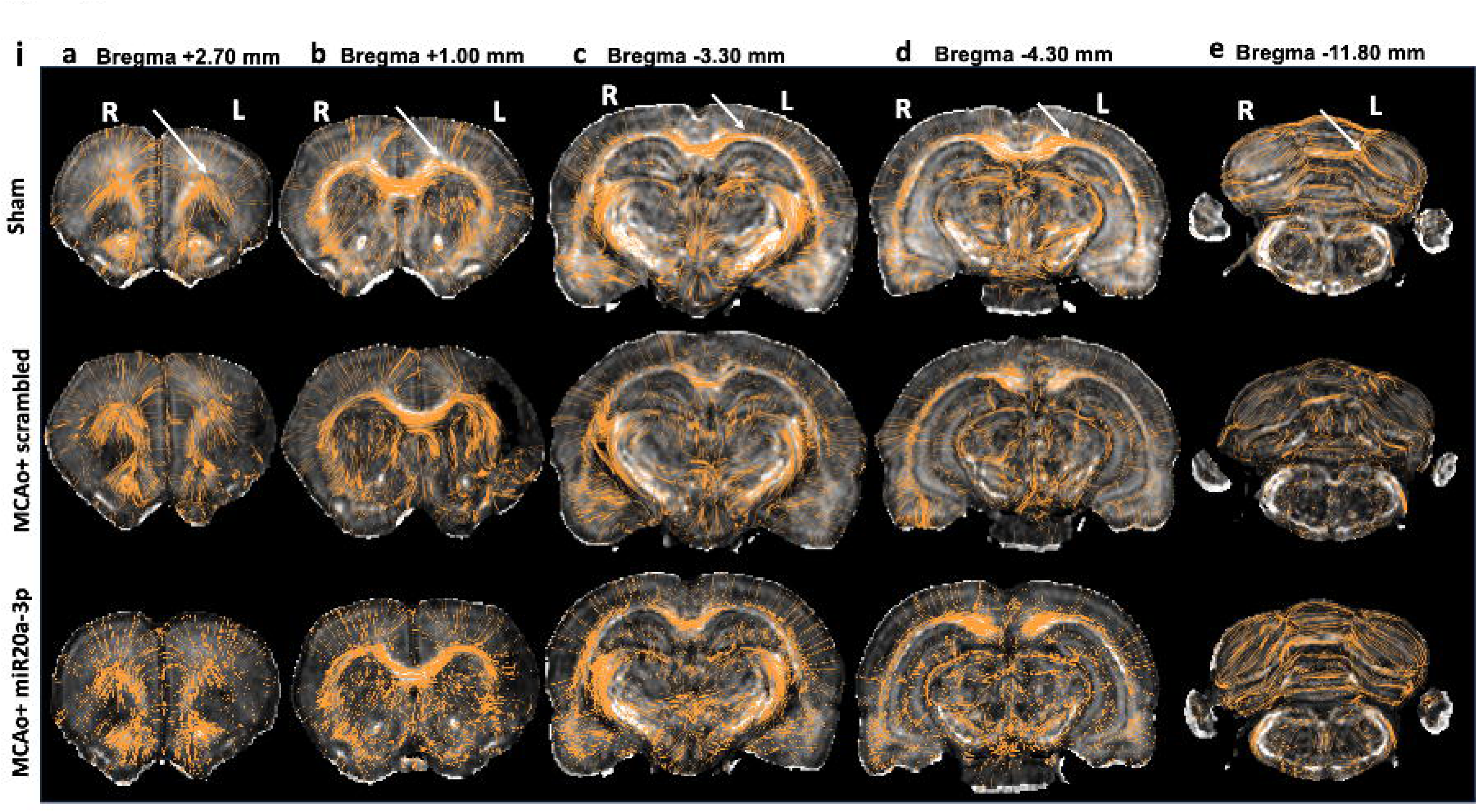
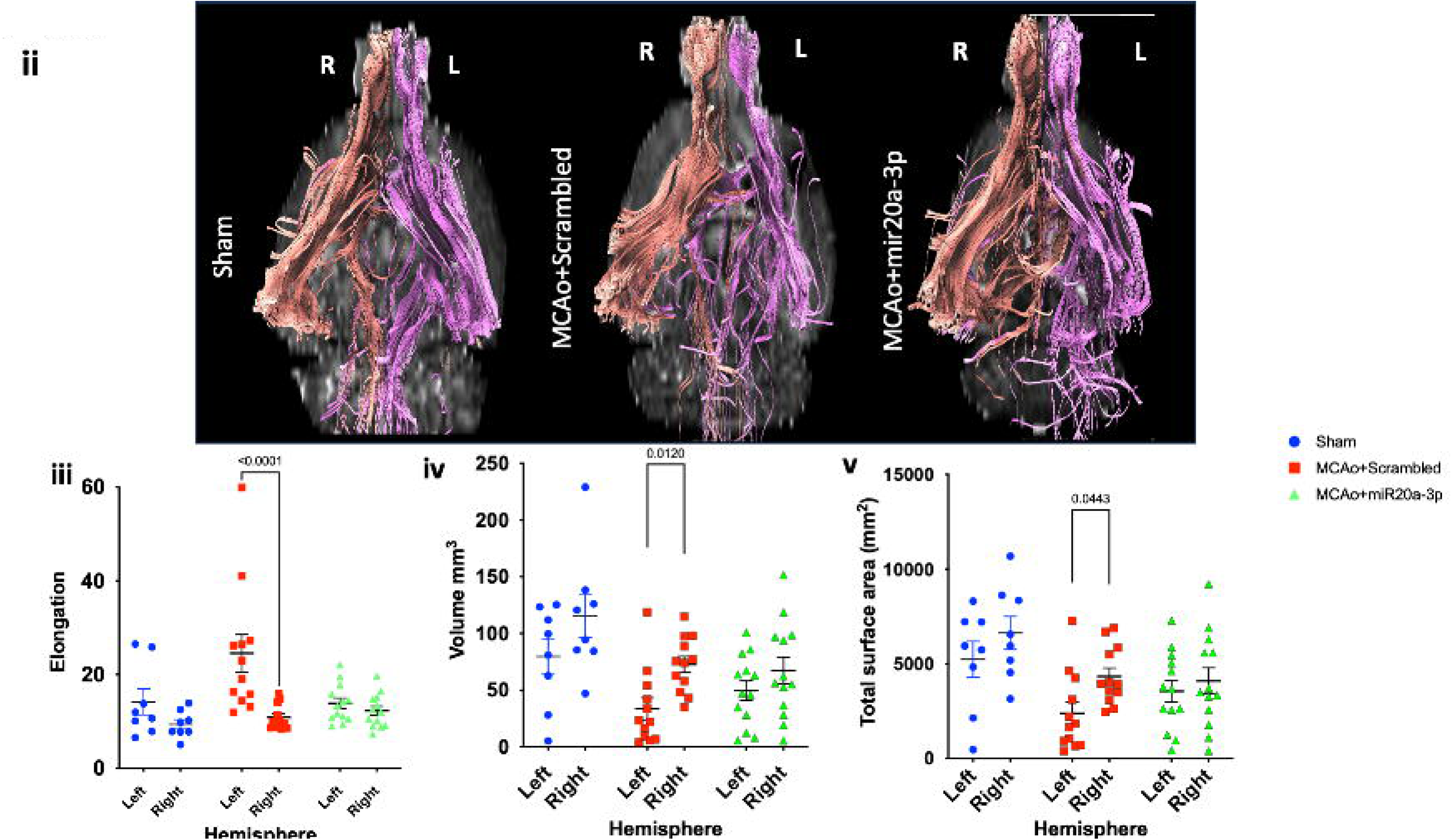
Altered connectivity due to stroke and mir20a-3p treatment: (i) Representative FA-DT MRI images from Sham, MCAo+Scr, MCAo+mir-20a-3p groups along the rostro-caudal extent of the brain. Location of each coronal image is indicated relative to Bregma (ii) Ventral view of the tracts between the anterior commissure (AC) and the basolateral nucleus of the amygdala (BLA), and their interhemispheric connections. R: right hemisphere, L: left hemisphere, tracts depicted in pink gold on the left hemisphere and rose gold on the right hemisphere. (iii-v) In each hemisphere of Sham, MCAo+Scr, MCAo+mir-20a-3p groups, AC-BLA tracts were assessed for (iii) Elongation, (iv) Volume (mm^3^) and (v) Total surface area (mm^2^). Left hemisphere is the ischemic hemisphere, right hemisphere is the non-ischemic hemisphere. N= 8 in sham, 12 in MCAo+Scrambled (MCAo+Scr), and 13 in MCAo+ miR-20a-3p groups. Significant p values are indicated on the histogram.

The anterior commissure (AC) connects the basolateral amygdala (BLA) across the two hemispheres and this connection is crucial for processing emotions, complex cognitive functions like fear memory, and aspects of decision-making. For the delineation of this connection, seed regions were placed in the AC and in the BLA for reconstruction. Evaluation of tract shape was executed with DSI. Ventral views of the reconstructed tracts between the AC and BLA are shown in Fig 2ii, on the left (ischemic; pink gold) and right (non-ischemic; rose gold) hemispheres. The tract reconstruction shows sparse tracts in the ischemic (left) hemisphere of an MCAo+Scr animal, but relatively similar tract densities in the Sham and MCAo+Mir-20a-3p animals. Quantification of shape analyses revealed that fiber elongation was significantly increased in ischemic hemispheric tracts as compared to non-ischemic hemispheres in the MCAo+Scrambled group (Fig 2iii), while tract elongation was similar between hemispheres of sham and MCAo+miR-20a-3p treated animals. Left-right asymmetry was also noted in tract volumes (Fig 2iv) and total surface area (Fig 2v) in the scrambled-treated stroke group, with significant reduction of tract volume and total surface area on the left (ischemic) hemisphere compared to the right (non-ischemic) hemisphere. No asymmetry was seen in the sham and miR-20a-3p treated stroke groups.

### Myelin staining reveals ischemic/non-ischemic hemisphere differences after stroke, which are ameliorated by miR-20a-3p treatment

In addition to MR imaging, long-term changes in brain morphometry were analyzed in Weil myelin-stained sections from the same animals. Histological sections were analyzed for hemisphere volume and ventricular volume as a gross measure of neurodegeneration as well as for major forebrain fiber tracts. The volume of the left (ischemic) hemisphere was significantly reduced in both stroke groups as compared to the right (non-ischemic hemisphere) while the two hemispheres were no different in Shams (Fig 3A). Lateral ventricular volume was similar in the two hemispheres of the Sham and MCAo+Mir-20a-3p treated animals (Fig 3B) but enlarged lateral ventricle volumes, an indication of neurodegeneration, were noted in the ischemic hemisphere of MCAo+Scr treated animals.

**Figure 3.**
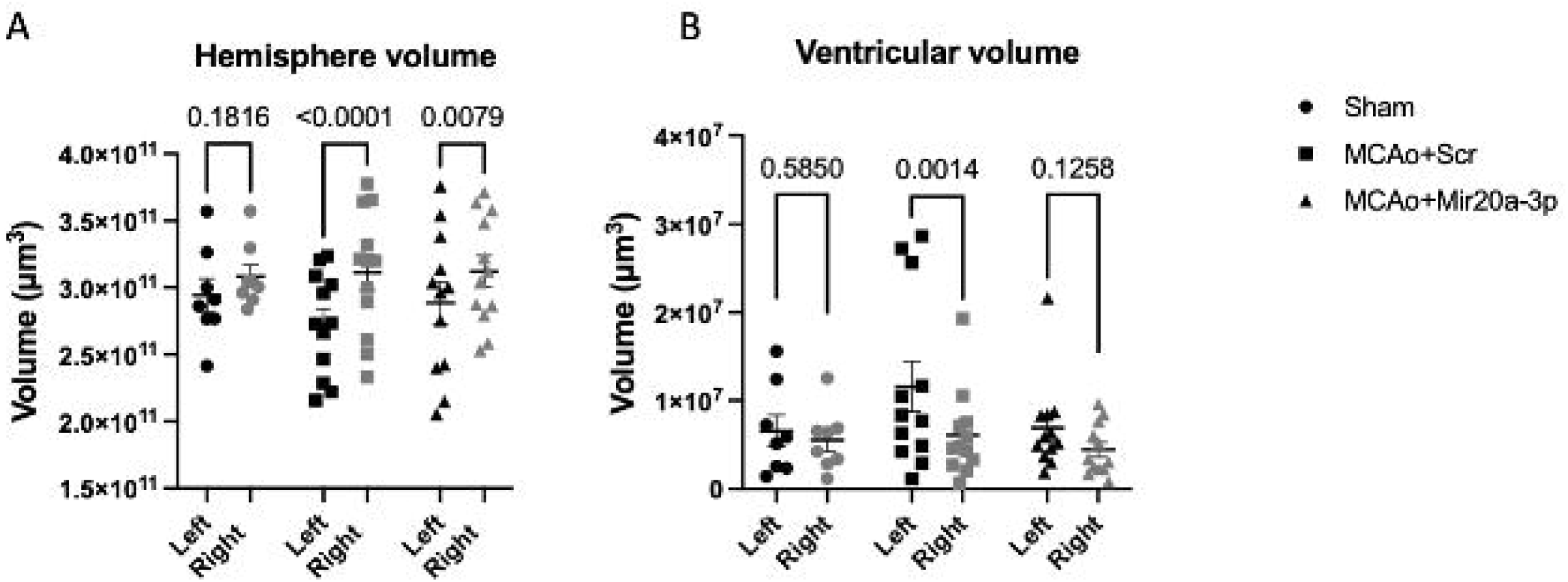
Effect of stroke and mir20a-3p on brain volumetric features (A) Volume (in um^3^) of the ischemic and non-ischemic hemisphere of Sham, MCAo+Scrambled and MCAo+mir20a-3p animals. (B) Lateral ventricular volume of the left (ischemic) and right (non-ischemic) hemispheres of Sham, MCAo+Scrambled and MCAo+mir20a-3p animals. N= 8 in sham, 12 in MCAo+ scrambled, and 12 in MCAo+ miR-20a-3p groups. Exact p values shown on the histogram represent comparisons of the left and right hemisphere. Key: black-filled shape: left (ischemic) hemisphere, gray-filled shape: right (non-ischemic) hemisphere.

#### White matter tracts

Three forebrain tracts were selected for analysis: corpus callosum, anterior commissure and internal capsule (Fig. 4). The volume of each myelinated tract was determined separately for the left (ischemic) and right (non-ischemic) hemisphere.

**Figure 4.**
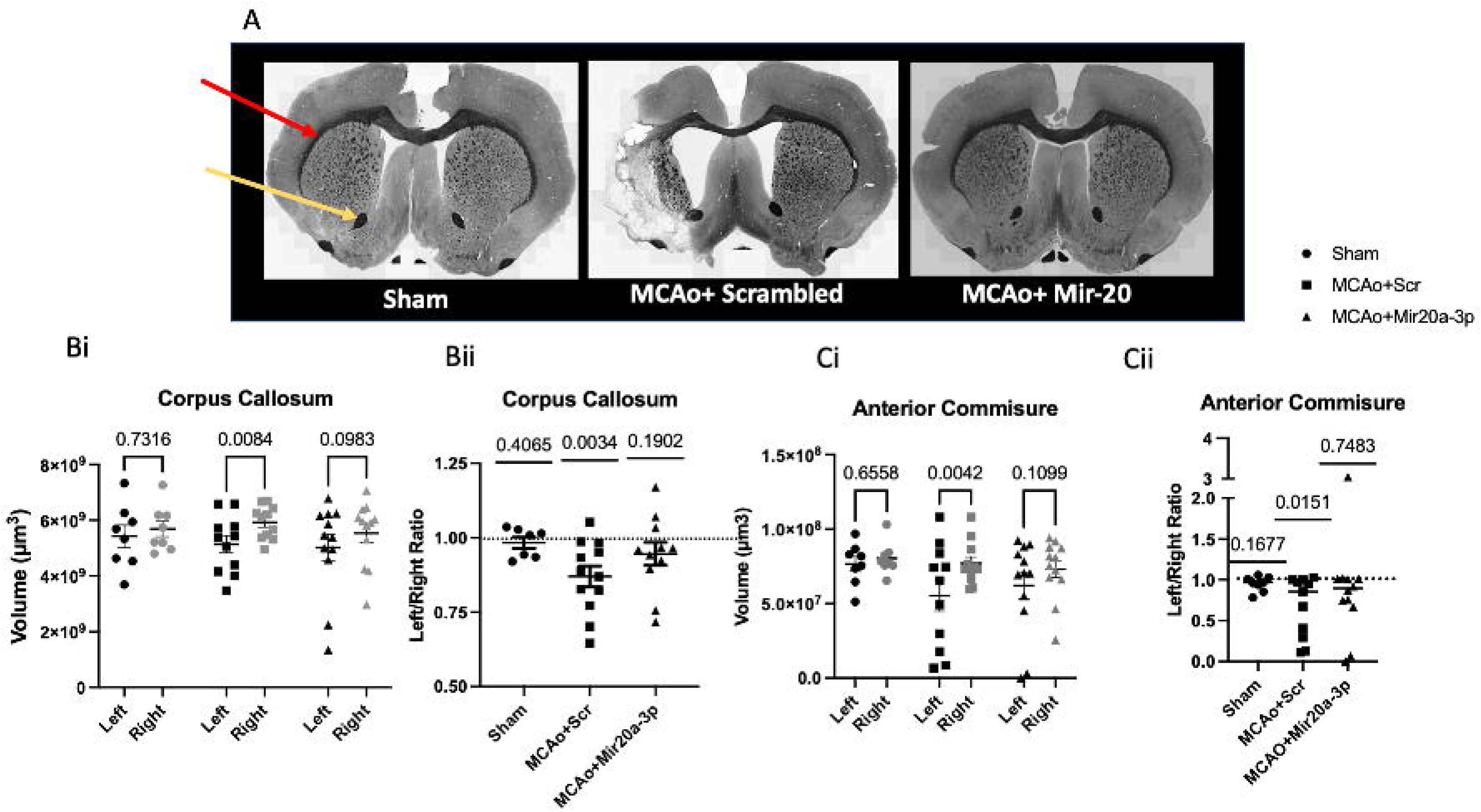
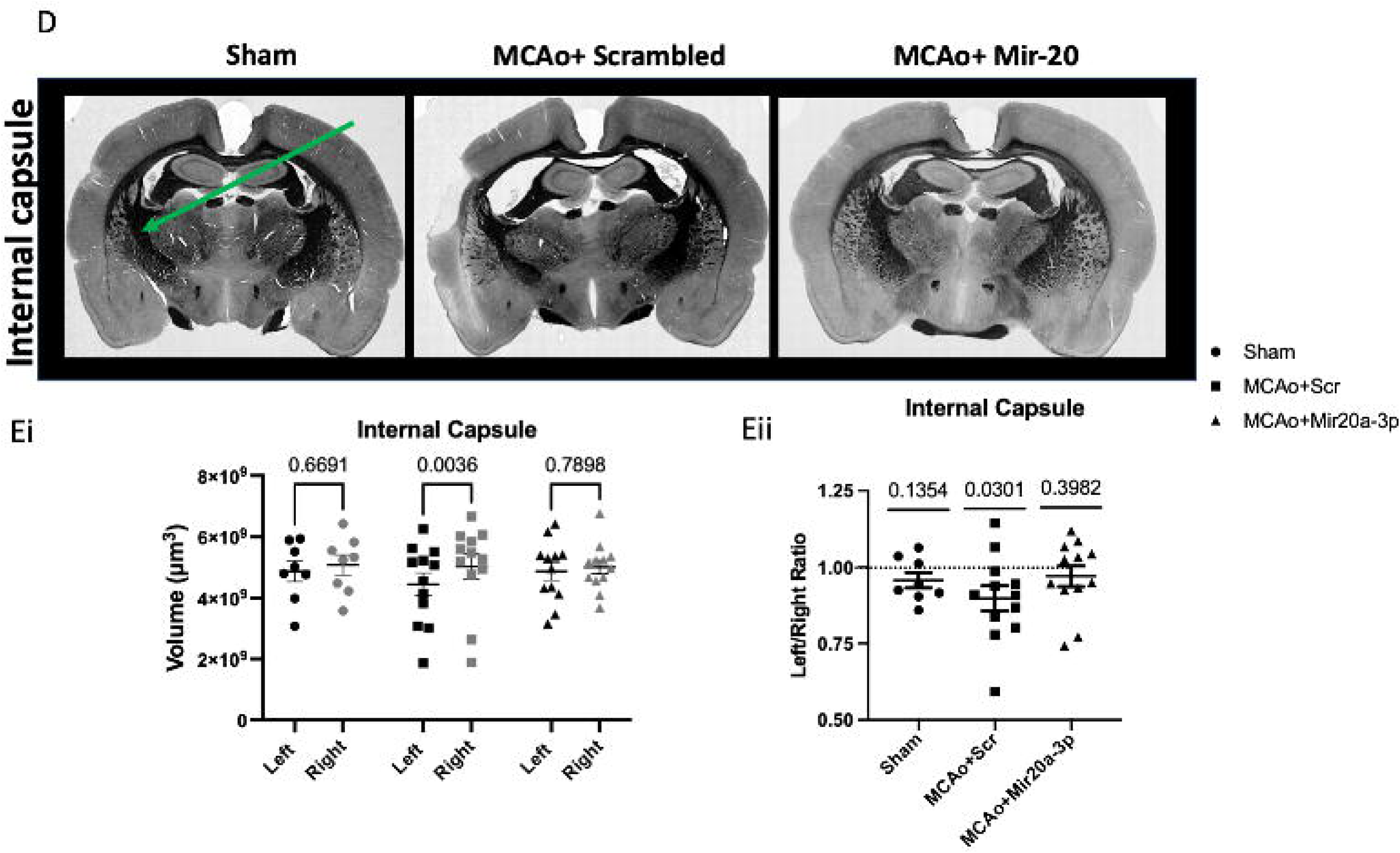
Volumetric changes in forebrain white matter tracts: White matter tract volumes were assessed in Weil myelin-stained sections. (A) Representative images of Weil myelin-stained rat brain sections at the cortico-striatal level. Corpus callosum indicated by red arrow, anterior commissure indicated by yellow arrow in the image from the Sham group. (B) Volume (um^3^) of the corpus callosum of the ischemic and non-ischemic hemisphere from Sham, MCAo+Scrambled and MCAo+mir20a-3p groups shown separately for each hemisphere (Bi) and ischemic side normalized to the non-ischemic hemisphere (Bii). (C) Volume (um^3^) of the anterior commissure from the ischemic and non-ischemic hemisphere from Sham, MCAo+Scrambled and MCAo+mir20a-3p groups shown separately for each hemisphere (Ci) and ischemic side normalized to the non-ischemic hemisphere (Cii). (D) Representative images of Weil myelin coronal sections at the level of the internal capsule, indicated by green arrow in the image from the Sham group. E) Volume (um^3^) of the internal capsule from the ischemic and non-ischemic hemisphere from Sham, MCAo+Scrambled and MCAo+mir20a-3p groups shown separately for each hemisphere (Ei) and ischemic side normalized to the non-ischemic hemisphere (Eii). Two-way ANOVA followed by paired comparisons (Bi, Ci, Ei), Wilcoxon one sample t-test (Bii, Cii, Eii) compared to the expected ratio of 1. Exact p values are shown on the graph. N= 8 in sham, 12 in MCAo+ scrambled, and 12 in MCAo+ miR-20a-3p.

#### Corpus Callosum

Representative images of the Weil myelin-stained sections from Sham, MCAo+Scr and MCAo+Mir-20a-3p groups depicting the corpus callosum (indicated by red arrow) and anterior commissure (indicated by yellow arrow) are shown in Fig 4A. The volume of the corpus callosum was similar in the left and right hemisphere of Sham animals but was decreased in the ischemic (left) hemisphere of the MCAo+Scr group as compared to the non-ischemic (right) hemisphere (Fig 4Bi). There was no difference in callosal volumes of the left and right hemispheres of the MCAo+Mir-20a-3p treated group (Fig 4Bi). Left/right callosal volume ratios were calculated and assessed by Wilcoxon one sample test to determine deviation from the expected ratio of 1 (ExpR-1). No significant differences from the ExpR-1 was seen in the Sham and MCAo+Mir-20a-3p group, but this ratio was significantly lower than 1 in the MCAo+Scr group (Fig 4Bii), indicating loss of myelinated fibers in the ischemic hemisphere.

#### Anterior commissure

Similar to the corpus callosum, the volume of the anterior commissure was similar in the left and right hemispheres in Sham and MCAo+Mir-20a-3p groups, while this tract volume was significantly reduced in the ischemic (left) hemisphere of the MCAo+Scr group compared to the non-ischemic hemisphere (Fig 4Ci). The ratio of left and right anterior commissure volumes were not different from the ExpR-1 in the Sham or MCAo+Mir-20a-3p groups but significantly lower than the ExpR-1 in the scrambled treated stroke group (MCAo+Scr) (Fig 4Cii) indicating loss of myelinated fibers in the ischemic hemisphere.

#### Internal capsule

Representative images from Sham, MCAo+Scr and MCAo+Mir-20a-3p groups depicting the internal capsule (indicated by green arrow) are shown in Fig 4D. Like the corpus callosum and anterior commissure, the volume of the internal capsule was comparable in the left and right hemisphere in the Sham and MCAo+Mir-20a-3p groups, while tract volumes were reduced in the left (ischemic) hemisphere in the MCAo+Scr group (Fig 4Ei). This group also showed a reduced left-right ratio from the ExpR-1 (Fig 4Eii). Overall, histological assessments of forebrain white matter tracts were consistent with FA DT-MRI findings.

### Stroke-induced microgliosis in the ischemic hemisphere is reduced by miR-20a-3p

#### Microglial activation in white matter tracts

Due to the reduced volumes of several forebrain tracts, we next examined the extent of microglia (Iba1), and myelin phagocytic microglia (presence of degraded myelin basic protein (dMBP) in Iba1+ cells). Single (Iba1+) and double-labeled (Iba1+dMBP) cells are shown in Fig 5Ai (enlarged view in Fig Aii). For subsequent analysis, the corpus callosum and the internal capsule were compared between scrambled and microRNA treated stroke groups.

**Fig 5.**
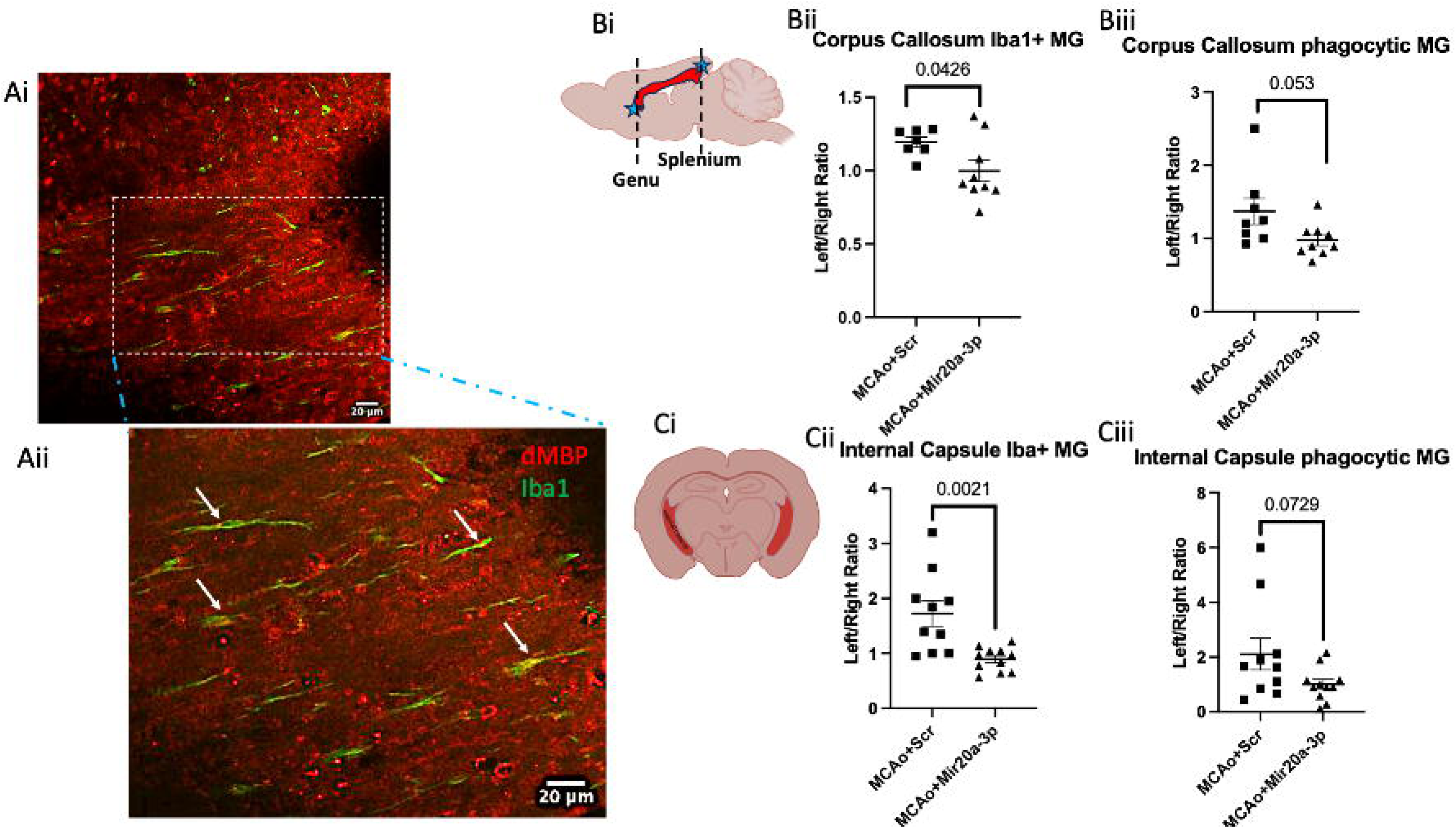
Abundance of microglia (Iba1+) and phagocytic microglia (IBA1+dMBP) in white matter tracts from stroke groups. (Ai) Representative image of Iba1+ microglial cells (green) and dMBP (red) immunohistochemistry from the genu of the corpus callosum. Region indicated by dashed white line is magnified and shown in Aii. Arrows in Aii indicate dMPB colocalized to Iba1 cells. (B) Quantitation of Iba1 positive and Iba1+dMBP double labeled cells from the corpus callosum. Bi. Schematic diagram shows the corpus callosum (red) in sagittal section. Cells were quantified in coronal sections through the genu and splenium of the corpus callosum (indicated by dashed lines). (Bii) Ratio of the total number of microglia in the left (ischemic) hemisphere and right (non-ischemic) hemisphere from the genu and splenium of the corpus callosum. (Biii) Ratio of the number of Iba1-positive microglia that colocalized dMBP in the left (ischemic) hemisphere and right (non-ischemic) hemisphere from the genu and splenium of the corpus callosum. (C) Quantitation of Iba1 positive and Iba1+dMBP double-labeled cells from the internal capsule. Ci. Schematic diagram indicates internal capsule in red (Cii) Ratio of the total number of microglia in the left (ischemic) hemisphere and right (non-ischemic) hemisphere from the internal capsule. (Ciii) Ratio of the number of Iba1 positive microglia that colocalized dMBP in the left (ischemic) hemisphere and right (non-ischemic) hemisphere from the internal capsule. N= 7-9 for Corpus callosum, 10-11 for Internal capsule. Exact p values shown on the histogram.

#### Corpus callosum

Iba+ microglia and Iba1+ dMBP+ cells in the genu and splenium (location indicated in the schematic in Fig 5Bi) of the corpus callosum were counted separately for each hemisphere. Number of single and double-labeled cells in the ischemic hemisphere was normalized to the non-ischemic hemisphere. Microglia numbers were significantly elevated in the MCAo+Scr group as compared to the MCAo+Mir-20a-3p group (p=0.0426) (Fig 5Bii), while comparisons of phagocytic microglia between MCAo+Scr group and MCAo+Mir-20a-3p showed a trend approaching statistical significance (p=0.053) (Fig 5Biii).

#### Internal capsule

Microglia and phagocytic microglia in the internal capsule (shown schematically in Fig 5Ci) were identified as described above. Microglia were significantly more abundant in the MCAo+Scr animals as compared MCAo+Mir-20a-3p group (p=0.0021) (Fig 5Cii) while comparisons of phagocytic microglia between MCAo+Scr group and MCAo+Mir-20a-3p showed a trend approaching statistical significance (p=0.0729) (Fig 5Ciii).

### Striatal microvessel density was increased by stroke and reduced by miR20a-3p

#### Tomato Lectin histochemistry

Sections from the cortex and striatum of Sham, MCAo+Scr, and MCAo+Mir-20a-3p animals were stained using fluorescently-labeled tomato lectin (Supplementary Fig S3). No group differences were seen in vessel density in the cortical region, while in the striatum, the MCAo+Scr group showed elevated vessel density compared to the MCAo+Mir20a-3 group (p<0.02).

#### Regression analysis

Regression analysis of histological measures related to white matter volume and microglial abundance were assessed for their contribution to behavioral impairment due to stroke. The following measures were included: L/R ratio of corpus callosum volume, internal capsule and anterior commissure, L/R ratio of microglia abundance in the corpus callosum and internal capsule, and L/R ratio of phagocytic microglia in the corpus callosum, and internal capsule. Coefficients, estimate and 95% confidence interval for each measure for the Novel Object Recognition Test and Remote Fear memory retrieval.

#### Novel Object Recognition Test

As described earlier, Sham, MCAo+Scr, MCAo+Mir-20a-3p were tested for episodic memory (NORT) pre stroke and at 30d and 100d days after stroke. At 100d, episodic memory, determined by the discrimination index, was not significantly different from pre-stroke values in Sham and MCAo+Mir-20a-3p group, but was significantly decreased in MCAo+Scr group, indicating impaired recall. Regression analysis showed that NORT was significantly associated with the volume of the corpus callosum (p=0.0172) and the abundance of microglia in this tract (p=0.115), as well as the abundance of microglia (p=0.002) and phagocytic microglia (p=0.0133) in the internal capsule. 95% confidence interval and p values are shown in Table 2A.

**TABLE 2A.**
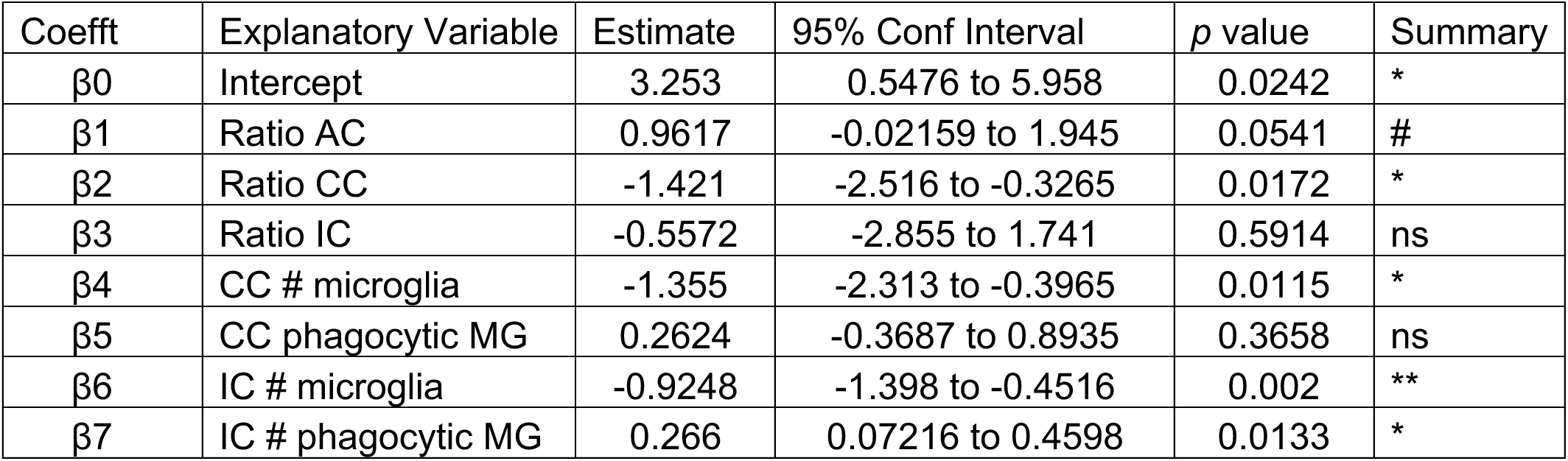
Estimates, confidence intervals, and p values for the linear regression models for Novel Object Recognition Test.

| Coefft | Explanatory Variable | Estimate | 95% Conf Interval | *p* value | Summary |
| --- | --- | --- | --- | --- | --- |
| $\beta_0$ | Intercept | 3.253 | 0.5476 to 5.958 | 0.0242 | * |
| $\beta_1$ | Ratio AC | 0.9617 | -0.02159 to 1.945 | 0.0541 | # |
| $\beta_2$ | Ratio CC | -1.421 | -2.516 to -0.3265 | 0.0172 | * |
| $\beta_3$ | Ratio IC | -0.5572 | -2.855 to 1.741 | 0.5914 | ns |
| $\beta_4$ | CC # microglia | -1.355 | -2.313 to -0.3965 | 0.0115 | * |
| $\beta_5$ | CC phagocytic MG | 0.2624 | -0.3687 to 0.8935 | 0.3658 | ns |
| $\beta_6$ | IC # microglia | -0.9248 | -1.398 to -0.4516 | 0.002 | ** |
| $\beta_7$ | IC # phagocytic MG | 0.266 | 0.07216 to 0.4598 | 0.0133 | * |

#### Remote Fear memory retrieval

Regression modeling for remote fear memory retrieval was also assessed, determined by the freezing response (Fig S2). Remote Fear memory retrieval was impaired early in the MCAo+Scr group (30 and 60d post stroke) but by 100d all groups showed poor recall on this task. Not surprisingly, none of these histological features were significantly associated with remote fear memory retrieval (Table 2B).

**TABLE 2B.** Estimates, confidence intervals, and p values for the linear regression models for Remote Fear memory retrieval.

| Coefft | Explanatory Variable | Estimate | 95% Conf Interval | *p* value | Summary |
| --- | --- | --- | --- | --- | --- |
| $\beta_0$ | Intercept | 154 | -61.74 to 369.7 | 0.1384 | ns |
| $\beta_1$ | Ratio AC | 69.95 | -8.465 to 148.4 | 0.0737 | ns |
| $\beta_2$ | Ratio CC | -40.67 | -128.0 to 46.64 | 0.314 | ns |
| $\beta_3$ | Ratio IC | -111.9 | -295.2 to 71.36 | 0.1968 | ns |
| $\beta_4$ | CC # microglia | 28.42 | -47.98 to 104.8 | 0.4159 | ns |
| $\beta_5$ | CC phagocytic MG | -49.69 | -100.0 to 0.6434 | 0.0524 | ns |
| $\beta_6$ | IC # microglia | 9.397 | -28.34 to 47.13 | 0.5816 | ns |
| $\beta_7$ | IC # phagocytic MG | -5.853 | -21.31 to 9.603 | 0.408 | ns |

## Discussion

We previously reported that middle-aged female rats exhibited impairments in associative and episodic learning in the chronic phase (30-100d) after ischemic stroke, while treatment with the small non-coding microRNA miR-20a-3p attenuated these cognitive deficits^21^. Since both cognitive tasks (fear conditioning and NORT) are mediated by circuits distal from the original cortico-striatal infarct site, the current study evaluated morphological changes in the brain in this behaviorally characterized female cohort, using a combination of DT-MRI imaging and histochemistry.

Our data showed that while there is ventricular enlargement and a gross reduction in hemisphere volume in the long term after stroke, ischemic injury preferentially impacts forebrain white matter tracts. We identified distinct group separations based on fractional anisotropy in white matter tracts between MCAo+Scr and MCAo+Mir-20a-3p groups. We also report on AC to BLA tractography differences, in tract shape characteristics between the left (ischemic) and right (non-ischemic) hemispheres. This was confirmed by histomorphometry analysis of Weil myelin-stained sections where left-right asymmetries were observed in the volume of the corpus callosum, internal capsule, and anterior commissure in the MCAo+Scr group, while treatment with miR-20a-3p mitigated this asymmetry. These data are consistent with reports that white matter hyperintensities on MR imaging serve as key biomarkers in stroke and vascular dementia, persisting beyond 90 days post-event^31^. These changes also correspond with other reports of neurodegeneration and related features of vascular dementia^31^. Similar studies by Yang et al^15^ have reported that DTI measures are affected after MCAo-induced ischemic stroke and correlate significantly with motor deficits and cognitive tests. In a long-term study of control and stroke patients, total brain volume by MRI and cognitive assessments across several domains showed greater total brain atrophy among stroke patients who displayed early cognitive impairment after stroke^32^.

Shrinkage of white matter tracts may reflect an overall pattern of neurodegeneration, initiated at the outset by the loss of neurons and their associated axons in the ischemic region. Decreased ischemic hemispheric volume in both MCAo groups and increased ventricular volume in the MCAo+Scr group suggest there is a loss of both white and gray matter. However, it is also possible that reduction in white matter volume may be due to loss of myelin resulting from ongoing persistent local inflammation. Elevated numbers of microglia and myelin phagocytic microglia were detected in the corpus callosum of the MCAO+Scr group compared to the MCAo+Mir-20a-3p group, suggesting the potential for a progressive loss of myelin even in the chronic phase of stroke. Pinter and colleagues ^33^ reported that activated microglia were present in the thalamus 7 months after embolic stroke, confirming long-lasting inflammation at sites remote from the initial lesion site. In addition to phagocytosis of myelin, loss of myelination may also be associated with impaired proliferation of oligodendrocytes (myelin-producing cells). Some support for this hypothesis comes from data showing that the mir17-92 cluster, containing mir-20a-3p, is highly expressed in oligodendrocytes and deletion of this cluster depletes oligodendrocytes^34^.

Our current study also underscores the therapeutic potential of microRNAs (miRNAs) as stroke treatments^35^. An extensive review highlights that several miRNAs are regulated following ischemic and hemorrhagic stroke therapy^35^, including miR-19a, miR-148, and miR-122 in patient populations^36^. In preclinical studies, miRNA have been shown to modify stroke outcomes, such as miR-1 and let-7f through regulation of IGF-1^37^; miRNA miR-29b, which prevents blood-brain barrier disruption^38^, miR-424^39^ and miR-let-7c-5p^39, 40^ which suppress microglial activation. Lacking are studies that have examined the long-term impact of miRNA treatment on stroke outcomes. We previously showed that miR-363-3p improved acute outcomes in females but not in males and alleviated post-stroke depression and cognitive deficits in middle-aged female rats ^24–26^. Subsequently, other studies have shown age- and sex-dependent expression of miRNAs, such as miR-15a, miR-19b, miR-32, miR-136, and miR-199a-3p, which are involved in key signaling pathways related to growth factors and cellular structure^23^. Mir-20a-3p, the focus of this study, was identified through miRNA profiling of post-stroke astrocytes and was significantly elevated in young animals compared to middle-aged females. Mir20a-3p mimetics injected iv to middle-aged females after stroke improved both acute ^20^ and long-term^21^ stroke outcomes. Mir-20a-3p is part of the miR-17-92 cluster, which regulates hippocampal neurogenesis^41^, influences anxiety and depression behaviors ^42^, and enhances neuroplasticity^43^. Additionally, miR-20a-3p promotes oligodendrocyte differentiation and neurite extension via mechanisms involving HspB1 and Rho signaling pathways^44^.

Several studies, including the present one, suggest that microglial activation persists long after the ischemic event and may correlate with adverse stroke outcomes. Repeated positron-emisson tomography (PET) studies using [^11^C]PK11195 in an embolic stroke model in rats demonstrated that signs of neuroinflammation were virtually absent at the lesion site after a few months, while microglial activation was later detected at a site distal (thalamus) to the infarcted area ^17^. In a study of diabetic rats, MCAo exacerbated the development of post-stroke cognitive impairment, which was attenuated by treatment with the C21, an angiotensin II type 2 receptor agonist, through modulation of microglial polarization^18^. Similarly, silencing microglia through viral-mediated knockdown of the colony-stimulating factor 1 receptor CSF1R) prevented stroke-induced cognitive impairment in diabetic mice^19^.

A key goal of this study was to assess whether morphological changes in white matter tracts are associated with cognitive impairment. This was found to be the case in the episodic memory test (NORT), which was associated with several features such as corpus callosum volume, and the presence of activated/phagocytic microglia in select tracts. None of these features were predictive for fear conditioning, where strong freezing responses were seen at baseline but reduced in all groups at 100d after stroke. It is worth noting that the freezing response was already weakened in the MCAo+Scr group as early as day 30 after stroke, indicating that stroke may accelerate the decline in associative memory. It is likely that periodic histomorphometry assessment initiated earlier after stroke would show that white matter losses occur more rapidly in the MCAo+Scr group.

White matter volumes were preserved in the mir20a-3p group, with region-specific reductions in abundance and phagocytic microglia. While the mechanism(s) underlying this action is not known, miR-20a-3p regulates genes associated with inflammation such as IL-17A and MMP9 in the acute phase. Since mir20a-3p treatment was administered in the acute phase after MCAo it suggests that the beneficial effects observed in this study, may be due to miR20a-3p acting on early events to limit white matter damage. IL-17A is known to activate microglia ^45;46^ and IL17A and MMPs also act to weaken the blood brain barrier (BBB) ^47, 48^ ^49^. Thus mir20a-3p suppression of these molecules may not only suppress local activation of microglia but also limit trafficking of macrophages and T cells into the brain. We observed a trend towards decreased serum GFAP levels, a surrogate marker for BBB damage, in mir20a-3p treated rats^20^, which partially supports this hypothesis. Relatedly, a recent study reported that miR-20a is poorly expressed in peripheral blood samples in patients with MS^50^, a demyelinating disease associated with T cell dysregulation^51^. T cell and/or macrophage/microglial activation may promote angiogenesis after ischemia^52^. For example, increases in brain microvessel density near the ischemic margin was associated with increased numbers of macrophages ^53–55^. Increased vessel density, which has been noted in mouse ^56^ and postmortem human stroke samples^57^, was also seen in the ischemic striatum of scramble-treated stroke animals, but not mir20-3p treated animals.

Overall, this study shows that stroke results in forebrain hemispheric asymmetries in the chronic phase and suggests that miR-20a-3p may target local inflammation to preserve myelination and white matter tracts, potentially reducing cognitive decline after stroke. Our previous work shows that enriching neuronal or astrocytic miR-20a-3p in middle-aged female rats is neuroprotective in the acute phase. Further studies are needed to assess whether this approach results in long term neurobehavioral improvement and to uncover the precise cellular and molecular mechanisms underlying the actions of this microRNA.

## Acknowledgements

Supported by RF-AG042189 to FS, an Alzheimer’s Association Research Fellowship (Award #: AARF-21-849749) to DS. The authors acknowledge the assistance of the Integrated Microscopy and Imaging Laboratory at the Texas A&M College of Medicine. RRID:SCR_02163.

## Supplementary figures

**Figure S1:** Schematic representation of the experimental strategy

**Figure S2.** Cognitive assessment: Morphological assessments were obtained from a cognitively characterized cohort of females (from Sampath et al 2023). A. Test of episodic memory (novel object recognition test) was assessed pre-stroke, 30 and 100 days after stroke. (B) Remote fear memory retrieval was assessed pre, 30-, 60-, and 100-days post stroke. Two-way repeated measures ANOVA followed by Tukey’s posthoc test. For NORT: Sham N = 7, MCAo + scrambled N = 9, and MCAo + miR-20a-3p N = 9-10. For Remote Rear memory retrieval: Sham N = 8, MCAo + scrambled N = 12, and MCAo + miR-20a-3p N = 13. (Details in ^21^)

**Figure S3.** Lectin histochemistry of coronal sections from the cortex (A) and striatum (B) of the left (top row) and right (bottom row) hemisphere of Sham, MCAo+Scrambled and MCAo+mir-20a-3p groups. Ratio of vessel density between the left and right hemispheres of the cortex (C) and (D) striatum. One Way ANOVA followed by Tukey’s post hoc test. *: p<0.05.

