## Supplementary figures and images for "Loss of white matter tracts and persistent microglial activation in the chronic phase of ischemic stroke in female rats and the effect of miR-20a-3p treatment"

### Supplemental Figures S1-S3

#### Supplementary section

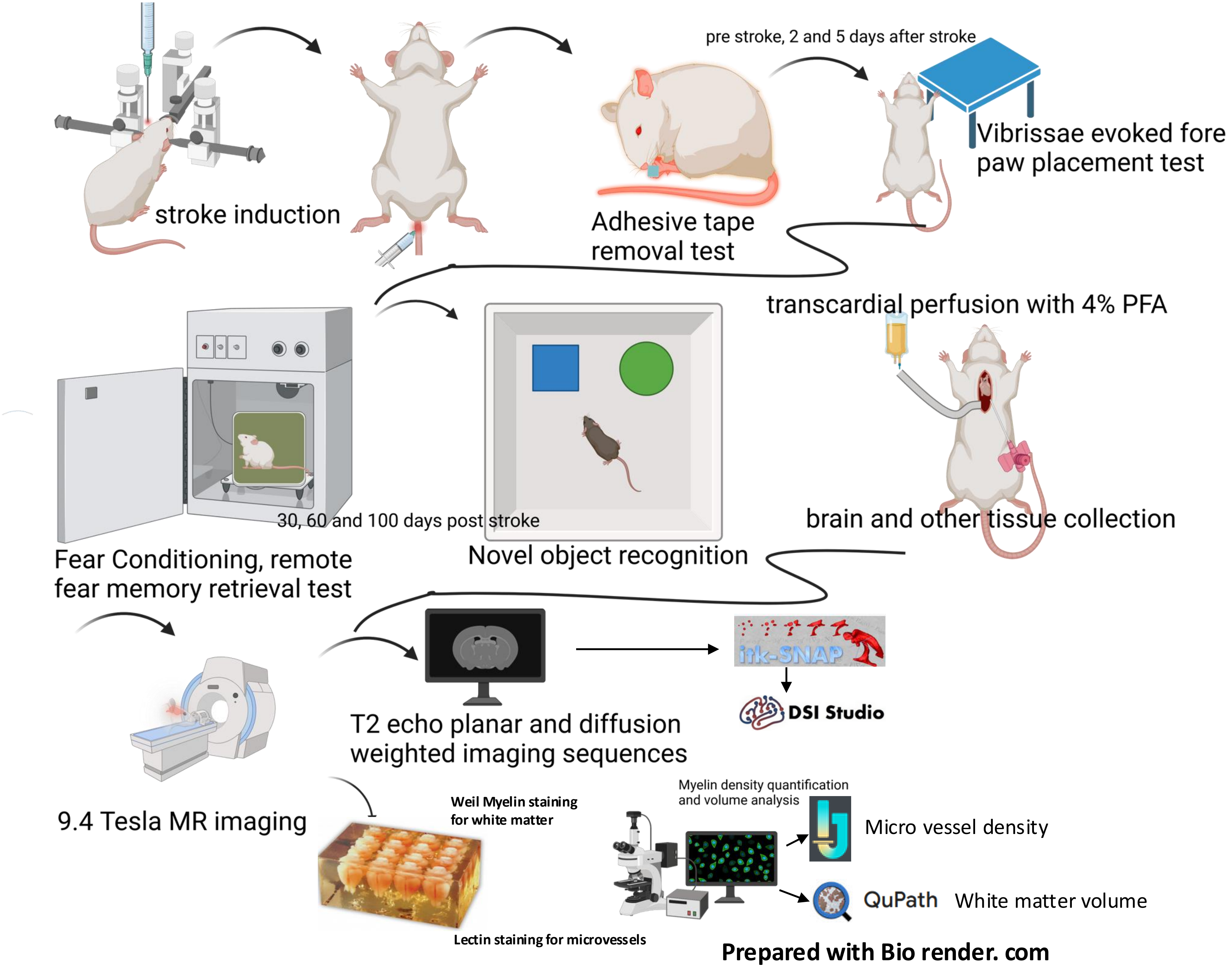

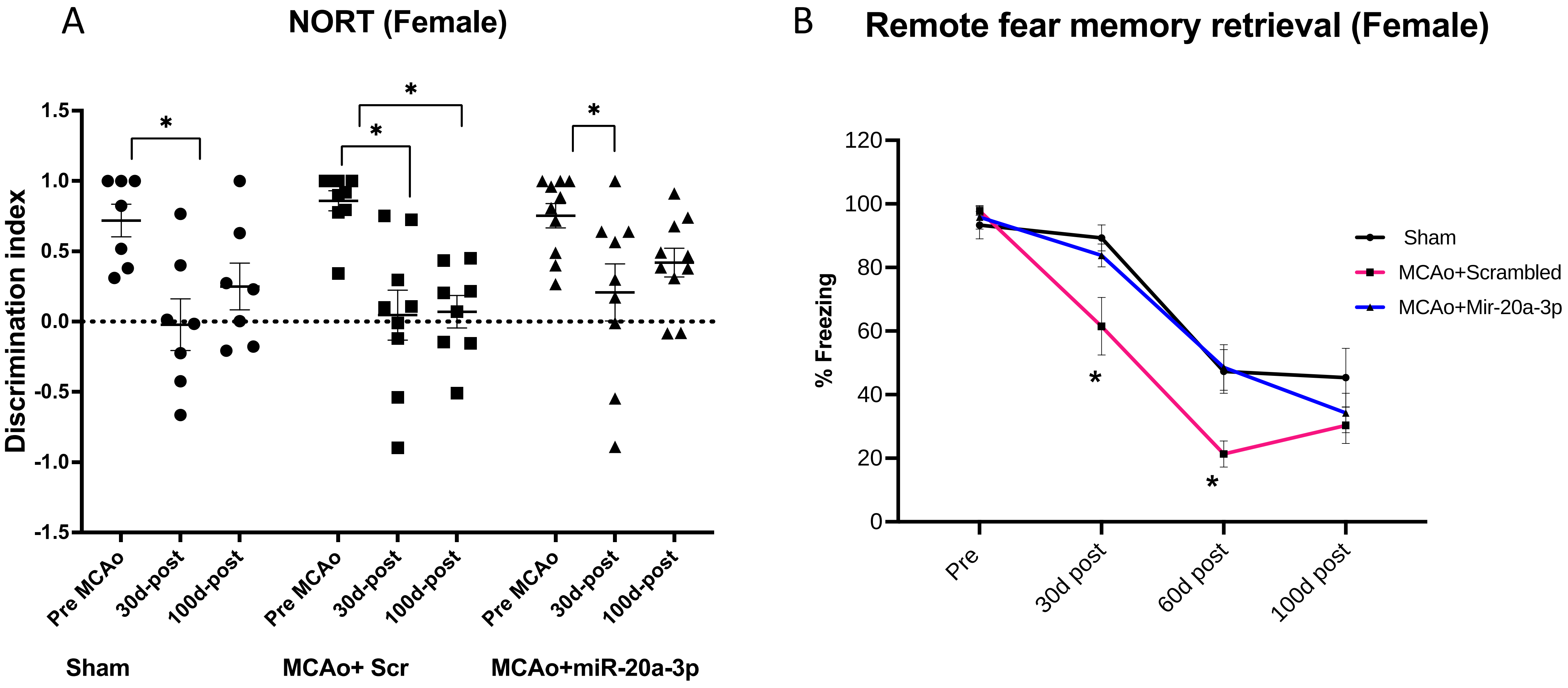

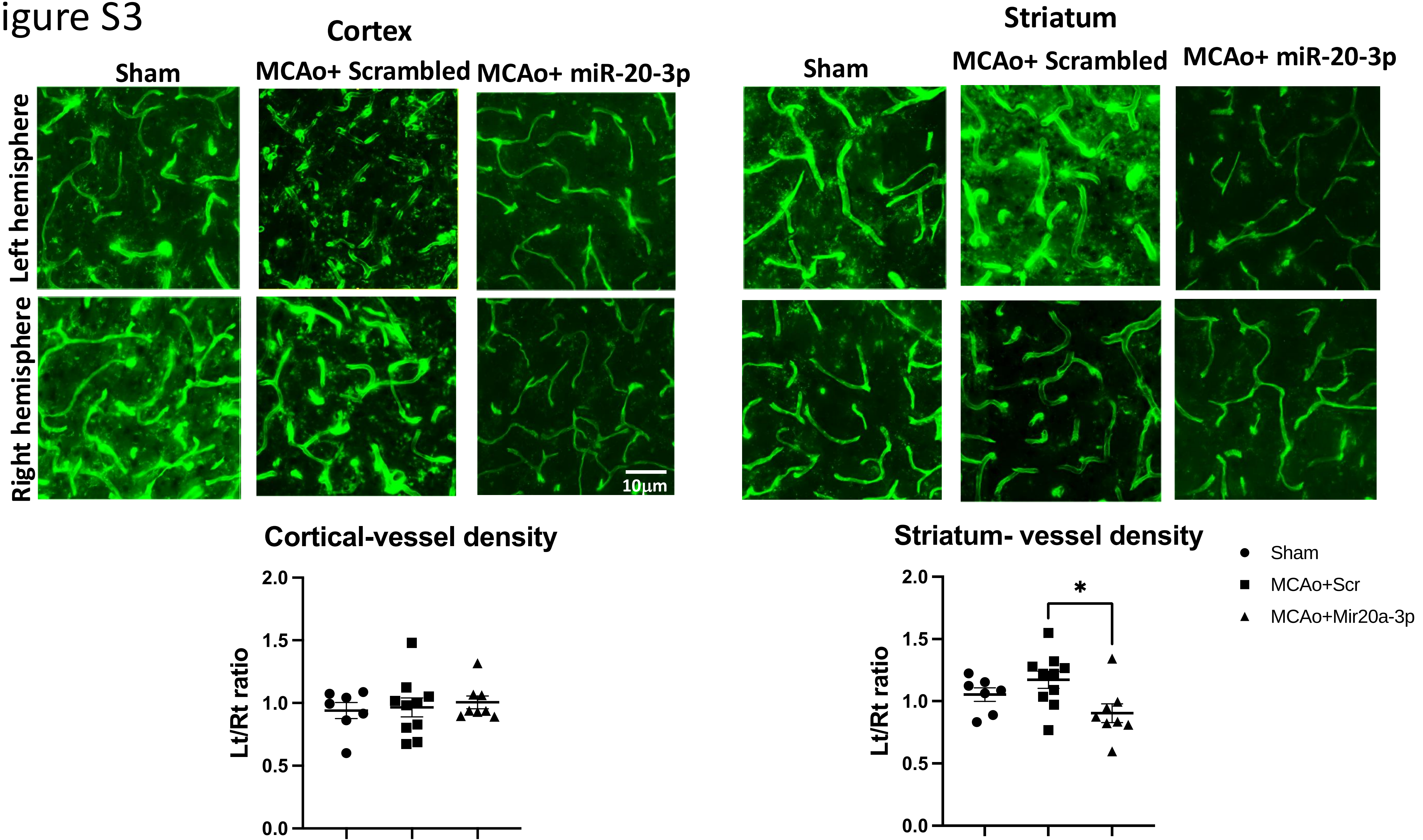
